# Neural evidence for the successor representation in choice evaluation

**DOI:** 10.1101/2021.08.29.458114

**Authors:** Evan M. Russek, Ida Momennejad, Matthew M. Botvinick, Samuel J. Gershman, Nathaniel D. Daw

## Abstract

Evaluating choices in multi-step tasks is thought to involve mentally simulating trajectories. Recent theories propose that the brain simplifies these laborious computations using temporal abstraction: storing actions’ consequences, collapsed over multiple timesteps (the Successor Representation; SR). Although predictive neural representations and, separately, behavioral errors (“slips of action”) consistent with this mechanism have been reported, it is unknown whether these neural representations support choices in a manner consistent with the SR. We addressed this question by using fMRI to measure predictive representations in a setting where the SR implies specific errors in multi-step expectancies and corresponding behavioral errors. By decoding measures of state predictions from sensory cortex during choice evaluation, we identified evidence that behavioral errors predicted by the SR are accompanied by predictive representations of upcoming task states reflecting SR predicted erroneous multi-step expectancies. These results provide neural evidence for the SR in choice evaluation and contribute toward a mechanistic understanding of flexible and inflexible decision making.

## Introduction

It has long been suggested that the brain can evaluate candidate actions by mental simulation, using internal maps or models of the environment (Tolman, 1948). For instance, even rodents can find novel shortcuts or flexibly plan new behaviors when their goals change (Dickinson, 1985; Tolman, 1948). Recent research drawing on model-based planning algorithms from reinforcement learning (RL), has offered a formal, computational framework for understanding how the brain accomplishes such behaviors (Daw, Niv, & Dayan, 2005; Doya, 1999; Keramati, Dezfouli, & Piray, 2011). These model-based planning computations involve enumerating, step by step, the different series of events (known as “states”) that might follow a candidate action, and evaluating the utility associated with each. But such exhaustive enumeration is infeasible in many tasks simply due to the large number of possible states (Daw & Dayan, 2014). Thus, despite the explanatory success of model-based planning as a framework for understanding behavior, the question remains open how the brain can actually implement some, necessarily simplified, version of these intractable computations.

For this reason, researchers both in artificial and biological intelligence have appealed to a variety of shortcuts for approximating planning computations using internal models. Most of these approaches ignore some information about the environment to simplify the model and reduce computational demand. However, by neglecting information that can be relevant to a choice, such simplifications necessarily lead to suboptimal choices, with potentially important implications. For instance, one maximally simplified evaluation scheme the brain is believed to use in some circumstances (known as model-free learning) leads to inflexible choices that may underlie phenomena of habits and also compulsive symptoms across several psychiatric disorders (Gillan, Kosinski, Whelan, Phelps, & Daw, 2016; Voon et al., 2015).

To achieve more flexible evaluation closer to full model-based planning while minimizing computational costs, other recent theories have suggested a role for *temporal abstraction*, such as chunking together multiple steps of a task (Botvinick, Niv, & Barto, 2009). One particularly appealing mechanism of this sort is the successor representation (SR) (Dayan, 1993; de Cothi et al., 2020; Geerts, Chersi, Stachenfeld, & Burgess, 2020; Madarasz & Behrens, 2019; Momennejad, 2020; Momennejad et al., 2017; Piray & Daw, 2019; Russek, Momennejad, Botvinick, Gershman, & Daw, 2017; Stachenfeld, Botvinick, & Gershman, 2017). SR reduces the computational costs of evaluation by storing predictions about future states, aggregated over multiple time-steps. Such aggregated state predictions can stand in for the step-by-step enumeration of an action’s long-run consequences, considerably simplifying later evaluation at the expense of potential failures when the environment changes in a way that invalidates the stored predictions.

There is suggestive evidence consistent with this predictive representation account, both from general expectancy-related neural activity (Barron et al., 2020; Boorman, Rajendran, O’Reilly, & Behrens, 2016; Bornstein & Daw, 2013; Brunec & Momennejad, 2020; Doll, Duncan, Simon, Shohamy, & Daw, 2015; Garvert, Dolan, & Behrens, 2017; Klein-Flügge, Barron, Brodersen, Dolan, & Behrens, 2013; Schapiro, Rogers, Cordova, Turk-Browne, & Botvinick, 2013; Stachenfeld et al., 2017; Wimmer & Büchel, 2019; Wise, Liu, Chowdhury, & Dolan, 2021) and more specific behavioral slips of action (Momennejad et al., 2017). But as yet, key tests of the mechanism have not been performed. In particular, it is not known whether anticipatory neural activity reflects the properties of the SR (e.g., that it spans multiple steps and neglects information the SR would lack). Furthermore, it is not known whether any such SR-aligned anticipatory neural activity preferentially accompanies the SR-predicted behavioral errors, when they occur.

A detectable and characteristic signature of the SR’s bundling of state predictions over multiple steps is predicted to occur when those aggregate summaries become inconsistent with knowledge of the individual step-by-step transitions. The SR for a state (the aggregate predictions of which states will follow it) is typically assumed to be learned directly from what follows the state when it occurs (Dayan, 1993). This makes it possible to provide experiences to a participant that establish expectancies about a series of states (e.g., states 1-3-5; see Fig. 1a), and then later expose the participant to a contingency change at a later step (e.g., 3-6) without providing the opportunity to directly observe its implications at an earlier step (i.e., that 1 will be followed by 6 rather than 5). The SR thus predicts that these manipulations should produce erroneous predictions about a state’s multi-step successors, and thus erroneous valuations of that state. Behaviorally, this prediction has been tested in “revaluation tasks” which measure whether participants are able to adjust their choices following such “transition revaluations”, in which the task’s state to state transitions are changed, as well as other analogous changes in contingencies. Here, across different types of change, the SR predicts a behavioral pattern of errors and correct responses distinct from fully model-based (and also model-free) approaches. We recently reported evidence that humans’ choices display a pattern of errors consistent with them sometimes computing value estimates using the SR’s stored multi-step state predictions (Momennejad et al., 2017). Importantly, participants did not always make these errors – implying that some of the time they successfully update their expectancies, e.g. via model-based evaluation or offline replay (Momennejad et al., 2017; Russek et al., 2017).

**Figure 1.**
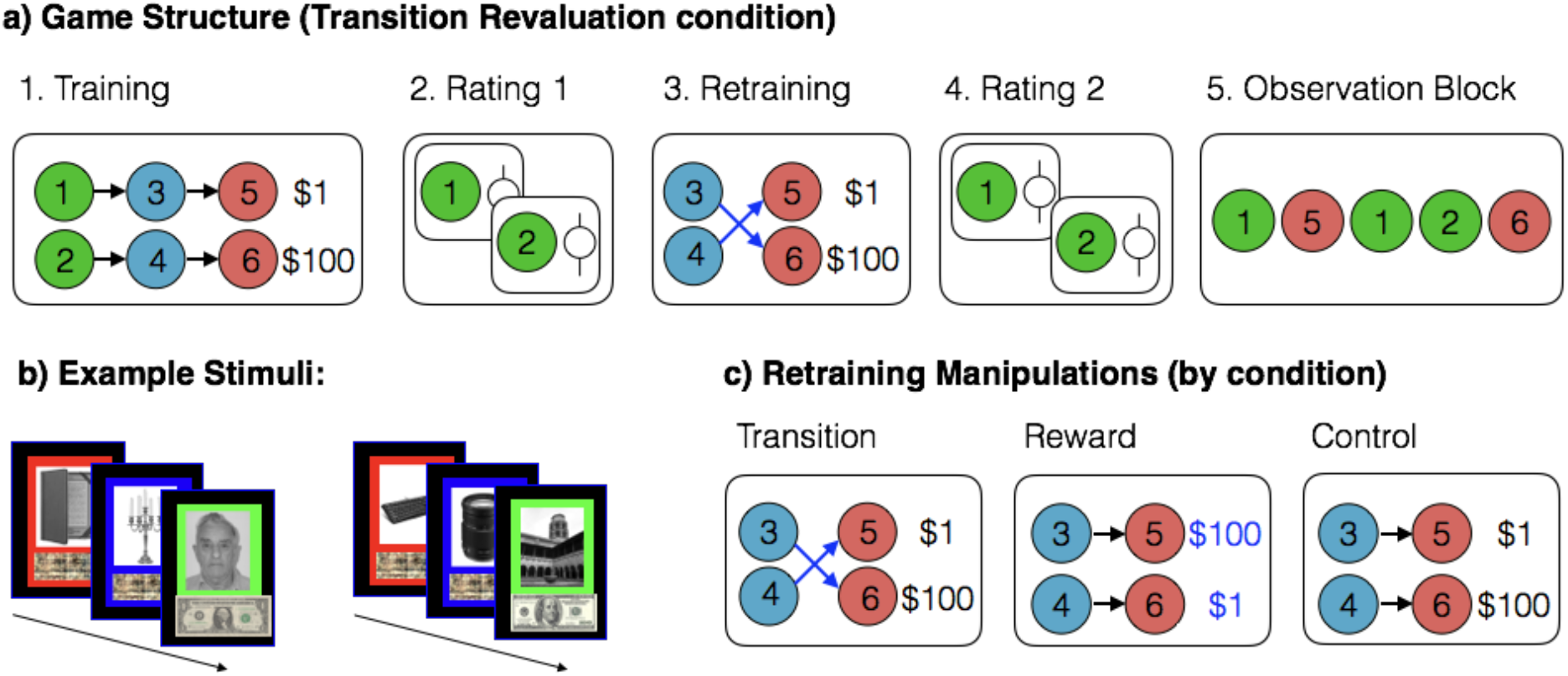
Game Structure. **a) Structure of states and rewards in a game from the Transition Revaluation condition.** Participants never saw these graphs and experienced the task structure one stimulus at a time. Each game consisted of 5 phases: Training, Rating 1, Retraining, Rating 2, and Observation Block. **b) Example visual images.** Each game had unique images; however, all games were arranged so that the first two states of either sequence were always object images, the last state of one sequence was a face image, and the last state of the other sequence was a scene image. **c) Retraining phase for each condition.** Every other phase was identical across conditions.

In the current study, we leverage this variability and use fMRI to investigate whether SR-predicted behavioral errors are indeed associated with neural representations of specific, SR-predicted, multi-step expectancies. In order to measure predictive representations during evaluation using fMRI, we used decodable categories of visual images to represent task states (Bornstein & Daw, 2013; Doll et al., 2015; Momennejad, Otto, Daw, & Norman, 2018). This allowed us to decode representations reflecting multi-step state predictions from prospective neural activity in the visual cortex during evaluation. By measuring multi-step predictions during evaluation, we were able to investigate whether participants’ behavioral errors suggestive of SR valuation were accompanied by decoded neural activity reflecting SR’s erroneous multi-step state predictions. We find evidence for this relationship, thus providing evidence for the SR as a process-level algorithm for choice evaluation in the brain.

## Results

### Behavioral Task

Forty human subjects underwent fMRI while performing an RL task designed to distinguish predictive neural representations and choices made by the SR from model-based (MB) and model-free (MF) strategies (Fig. 1; Momennejad et al., 2017). The task consisted of ten games. Each game began with a Training phase in which subjects were repeatedly exposed to two three-state sequences, composed of visual images, whose final states contained rewards of different magnitude (Fig. 1a). Images were organized so that the final state of either sequence (Fig. 1a, States 5 and 6) was from a different fMRI-decodable image category (either a face or a scene) that was also different from the category for the starting (States 1 and 2) and middle states (States 3 and 4; objects; Fig. 1b). In a Retraining phase, by experiencing a number of partial sequences starting from the middle state, subjects learned that either the final state to which each middle state transitioned (Transition Revaluation condition) or the reward in each final state (Reward Revaluation condition) had been switched, or that no changes to the sequences had been made (Control condition; Fig 1c). Before and after retraining, subjects successively viewed the first and middle images of either sequence, in random order, and rated their preference for starting a new sequence from that image.

The states and rewards were arranged so that, on each game, one of the two start states (State 2) would be initially preferable (i.e., leading to the larger reward) at Rating 1, but the Retraining phase in Transition Revaluation and Reward Revaluation games (but not Control games) should ideally reverse this preference, because the other start state (State 1) would now lead to the larger reward. SR makes behavioral predictions, distinct from MB and MF learning, about the pattern, across conditions, of errors in the valuation of the start images (i.e., the ability to correctly reverse preferences) during Rating 2. It also predicts that these errors follow from corresponding errors in predictive representations of the consequences expected to follow either start state. To measure such predictive representations, we decoded covert representations of the image categories for each final state, from fMRI data collected during Rating 2 for each start state. To localize where else in the brain such predictive activity occurs, we used RSA on fMRI data collected during an observation block, presented following the second rating of each image, during which subjects viewed 30 image presentations, consisting of the first and final state of each sequence, in random order (Fig. 1a; Supplementary Fig. 4).

### Pattern of behavioral errors reflects influence of SR

The key theoretical difference between the SR, and pure MB or MF algorithms is what they learn (i.e., in the different representations summarizing previous experience that they learn, update, and use to guide evaluation). Our behavioral and neural analyses are both based on detecting how those representations are affected by the retraining manipulation and therefore produce different patterns of correct or erroneous valuations during the subsequent, second rating. Importantly, these different RL strategies are not mutually exclusive; instead, much evidence suggests that the brain uses multiple decision strategies (corresponding, in this framework, to maintaining multiple representations of experience) and relies differentially on them across games. Accordingly, in practice, there is variation across both games and subjects in correctly solving such revaluation tasks (Barron et al., 2020; Gershman, Markman, & Otto, 2014; Momennejad et al., 2018, 2017; Wimmer & Shohamy, 2012), and we leverage this variation to test for associations between neural and behavioral markers of the SR strategy.

The qualitative signature of long-run, temporally abstract state predictions (as in the SR) that we seek to test is that these predictions can be rendered invalid by changes in subsequent state transition contingencies, which is detectable as a selective pattern of errors on games that probe this type of change (Transition Revaluation), relative to otherwise similar manipulations (Reward Revaluation). To motivate these analyses in more detail, we sketch the representations formed by each algorithm during initial training and how they change with retraining (Fig. 2a). The central representation used by MB learning is a “one-step” state transition model (i.e., for each state, which one comes next). During evaluation, predictions of states multiple steps into the future can then be made, iteratively, by chaining together a series of one-step predictions. In contrast, the SR learns and stores state predictions already aggregated over multiple steps: learning, for each state, the set of all subsequent states that will ultimately be visited by the end of the sequence. Both MB algorithms and SR also learn the immediate reward for each state (e.g. $100 in 6). Evaluation then consists of predicting states (either by iterative chaining, MB, or retrieving multistep predictions, SR) and adding up their associated rewards. Finally, MF algorithms do not predict states or one-step rewards at all, but rather store, for each state, the expected, long-run cumulative reward that will ultimately follow it.

**Figure 2.**
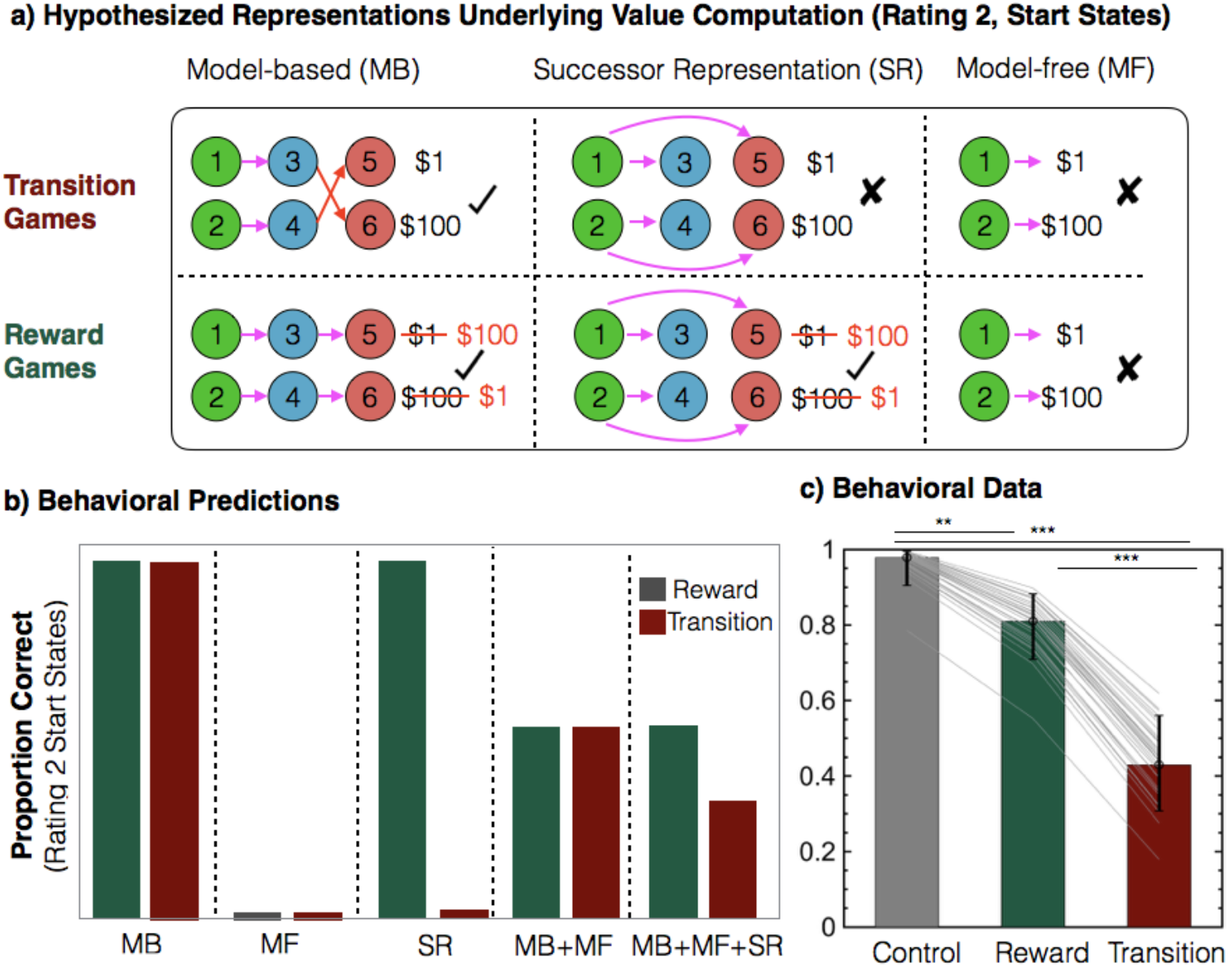
Hypothesized representations and behavioral results. **a) Representations learned by MB learning, SR and MF learning during Rating 2 for Transition Revaluation and Reward Revaluation conditions.** Note that arrows reflect representations formed by each algorithm, which may differ from the transitions experienced in the task. (Top) Transition Revaluation games. Because MB simulates final states using an updated one-step transition model whereas SR relies on multi-step predictions that are not updated, MB and SR predict that different final states will follow either start state. SR’s failure to predict the updated final state leads to an incorrect evaluation and a failure to reverse preference in Rating 2. (Bottom) Reward Revaluation games. Retraining causes no changes in which final states follow either start state. MB and SR predict the same final state and, unlike MF, both correctly revalue start states. (Top and Bottom) For both conditions, MF cannot update reward predictions without new experienced transitions from either start state. **b) Proportion correct Rating 2 responses for combinations of MB, MF and SR strategies across conditions.** Both SR and MB correctly alter valuations on Rating 2 following Reward Revaluation games. Only MB alters valuations on Transition Revaluation games. MF cannot update valuations following either condition. Since MB and MF succeed and fail respectively at both manipulations, combined in any proportion, they make errors for either condition in equal proportion. Combinations of strategies that include SR commit more errors in Transition Revaluation games than in Reward Revaluation games. **c) Measured Rating 2 responses reflect influence of SR in evaluation.** Proportion correct Rating 2 responses for start states (N = 39 subjects). Bar heights reflect measured proportion correct responses estimated from group-level, fixed effect estimates from a generalized linear mixed effects model relating Rating 2 correctness to condition. Error bars show 95% confidence intervals. Grey lines show fitted subject-level random effects estimates. **P < .01, ***P < .001, generalized linear mixed effects model.

Standard models assume that each of the representations is learned directly from what is experienced following visits to the various states (Dayan, 1993; Sutton & Barto, 2017). Thus, in particular, the SR, MB, and MF algorithms all learn about states 1 and 2 (their multi- or one-step successors, or reward values, respectively) during initial training, but retraining only affects what they know about states 3-6: because the initial states are not re-experienced, their predictions are left unaltered (Fig. 2a). Importantly, this does not mean that the algorithms do not change their evaluations of expected reward from the start states following retraining. This is because, for MB and SR, the value of states 1 and 2 is computed by retrieving which states are predicted to follow, and then combining this with their one-step rewards. In particular (Fig. 2b), MB successfully alters its ratings equally well following both types of revaluation (for Transition Revaluation, because it uses new one-step transitions to discover the new final states; for Reward Revaluation by taking account of the new value of the final states). In contrast, SR succeeds at Reward Revaluation games (because it incorporates the new value of the old final state) but fails at Transition Revaluation games (because the now-erroneous multi-step prediction from the start state is not updated). Since MF learns only completed long-run reward predictions, the retraining phase has no effect in any condition, and it does not revalue.

Altogether, the predicted behavioral signature of SR is that revaluations for the starting state (at Rating 2) will be less frequently correct in Transition Revaluation games relative to Reward Revaluation games (Fig. 2b). Note that since MB and MF learning each perform equivalently for both conditions, no mixture of those strategies alone can explain a difference between conditions.

To test this behavioral prediction, we examined whether Rating 2 performance was worse on Transition Revaluation games than on Reward Revaluation games. Specifically, we measured change in a categorical measure of correct revaluation at Rating 2 (correct versus incorrect start state preference) as a function of condition (Reward Revaluation vs Transition Revaluation games). Since revaluation is only informative if learning during initial training and retraining succeeds, for all analysis we removed games in which learning was not successful during either training or retraining, as indicated by Rating 1 ratings for all states as well as Rating 2 ratings for middle states, as well as games for which subjects did not indicate clear preferences (Methods).

To measure differences in successful revaluation between conditions, we used a generalized linear mixed effects model predicting successful revaluation as a function of condition (Methods). This approach revealed that, consistent with our hypothesis, subjects correctly revalued start states less frequently on Transition Revaluation games compared to Reward Revaluation games (Fig. 2c; Contrast estimate = −1.735, F(1) = 23.704, P < .001), replicating our previous study (Momennejad et al., 2017). Subjects also had correct Rating 2 preferences more frequently on Control games compared to Reward Revaluation games (Contrast estimate = −2.3648, F(1) = 8.4789, P = .004) as well as on Control games compared to Transition Revaluation games (Contrast estimate = 4.0997, F(1) = 25.881, P < .001). This provides evidence that successful revaluation on Reward Revaluation and Transition Revaluation games was not caused by nonspecific factors such as forgetting which start state was followed by which final state during Training, which would lead to equivalent preference changes in control games as well. Finally, we note that these results are robust to defining revaluation success using a continuous measure (Revaluation Score; (Momennejad et al., 2017)), as well as to using other strategies for controlling for Retraining acquisition failures, which remove fewer games (Supplementary Fig. 1).

### Predictive neural activity in visual cortex relates to SR-predicted pattern of behavioral of errors

To further test the interpretation that this pattern of behavioral errors resulted from erroneous multi-step predictions, as reflected in neural representations, we used fMRI to assess state expectancy more directly. We asked whether neural signatures of expectancy tracked the SR’s multi-step state predictions, and, in particular, whether such a match co-occurred with the SR’s putative behavioral signature: erroneous post-retraining ratings on Transition Revaluation games (Fig. 3a).

**Figure 3.**
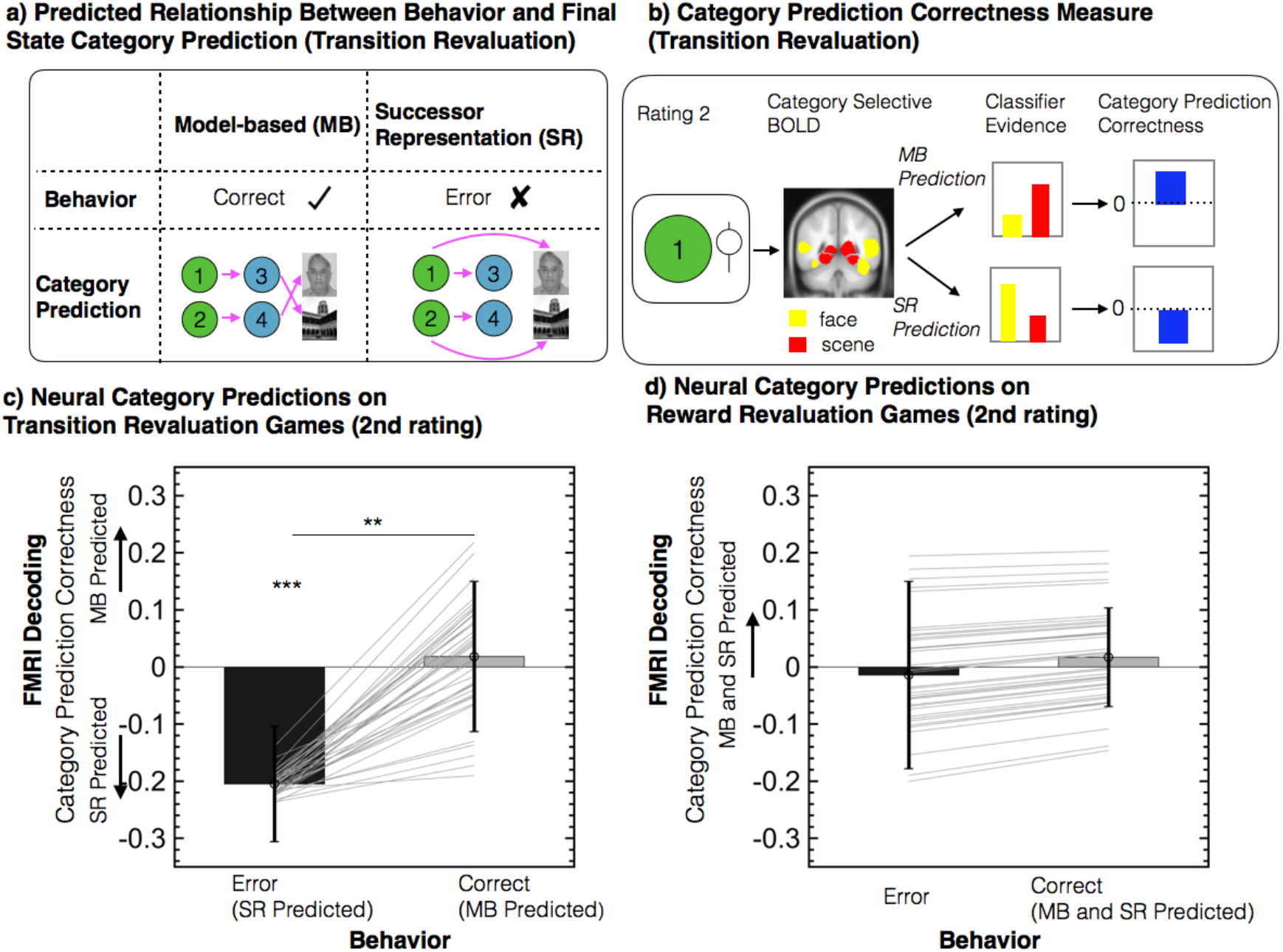
Decoding predictive representations during evaluation. **a) SR responsibility for behavioral errors predicts that erroneous ratings should be accompanied by erroneous final state category predictions.** Predicted relationship between behavior and final state category predictions, for the example Transition Revaluation game from Fig 1a. In this game, state 5 was a face image and state 6 was a scene image. During Rating 2, SR predicts the erroneous final state and produces erroneous valuations. In this example game, it predicts that incorrect ratings for Image 1 should be accompanied by Face category predictions. MB predicts the correct final state that will follow either start state and produces correct valuations. In this example game, it predicts that correct ratings for Image 1 should be accompanied by Scene category evidence. On Reward Revaluation games (not shown), because no Transition changes occur, both MB and SR predict the same final state image categories, as learned in Training, and both predict correct behavior. **b) Category Prediction Correctness measures extent to which category evidence reflects correct state predictions.** During Rating 2 for each start and middle state, BOLD data in face and scene category selective regions was input to classifier trained to classify viewing of faces, scenes or objects (note: rated state image was always an object). Category Prediction Correctness measures extent to which evidence from this classifier for faces versus scenes reflects the category of the state that would be correctly expected to follow the state being rated. On Transition revaluation games, MB predicts Category Prediction Correctness to be positive (e.g. Scene > Face when rating Image 1), SR predicts it to be negative (e.g. Scene < Face when rating Image 1). On Reward revaluation games (not shown), both MB and SR predict positive Category Prediction Correctness. **c) Transition Revaluation errors are accompanied by SR-aligned erroneous category predictions.** Measured Category Prediction Correctness on Transition Revaluation games, split by whether revaluation behavior was incorrect (consistent with SR) or correct (consistent with MB). **d) Category predictions do not change with behavior on Reward Revaluation games.** Measured Category Prediction Correctness on Reward Revaluation games, split by whether revaluation behavior was incorrect (consistent with MF) or correct (consistent with SR and MB). **c, d)** Bar heights show fixed effects group prediction from a linear mixed effects model predicting Category Prediction Correctness, applied to ratings from 39 subjects. Error bars show 95% confidence intervals. Grey lines show subject-level random effects predictions. *P < .05, ** P < .01, *** P < .001, linear mixed effects model.

To measure multi-step predictive representations, we used data from a visual localizer task (collected prior to the main task) and trained a logistic regression classifier to decode from multivariate BOLD activity in visual cortex whether a subject was viewing an image of a face, scene, or object (Methods). We then applied this classifier to measure covert neural evidence for faces versus scenes during Rating 2, while a subject evaluated start states, always objects. We treated this relative evidence as a read-out of the multi-step predictive state representation elicited during evaluation (Fig. 3b). For each game, we defined a score (“Category Prediction Correctness”) measuring the extent to which face versus scene evidence provided by the classifier, reflected the category (face or scene) of the final state that would correctly be expected to follow the state being rated, conditioned on correct integration of the Retraining phase information into the prediction. For each game, we computed this measure over the second rating for the start state of each sequence and averaged them.

Our primary interest was in examining whether behavioral responses suggestive of SR computations are driven by neural activity reflecting SR predictive representations. We thus examined Transition Revaluation games, where, during the second rating, MB and SR make opposite predictions about which final state will follow either start state. In particular, following the Transition change, MB arrives at newly correct multi-step predictions by re-simulating a one-step transition model. In contrast, SR re-uses the, now erroneous and outdated, multi-step transition predictions learned in the training phase during Rating 2. The key prediction for the hypothesis that SR behavioral responses are driven by SR predictive neural activity is thus that incorrect behavioral Transition revaluation responses will be accompanied by neural activity reflecting incorrect category predictions (Fig. 3a). We tested this prediction by examining Transition Revaluation games for which errors were committed. On these games, we observed that Category Prediction Correctness was negative (Estimate: −.219, F(1,61.154) = 5.118, P = .027), thus supporting the hypothesis that errors on Transition Revaluation games are driven by SR over MB predictions. This provides evidence for our main prediction, that behavioral errors predicted by SR evaluation are driven by neural activity reflecting SR state predictions.

We next examined whether SR-predicted erroneous neural activity occurred selectively when behavior reflected SR predictions. We thus examined Transition Revaluation games where subjects made correct revaluations, thus reflecting MB behavioral predictions, and whether, on successful performance, neural activity was aligned with MB predicted correct category predictions. On these games, although Category Prediction Correctness was not significantly positive (Estimate: 0.0184, F(1,26.921) = 0.0759, P = 0.785), it was less negative than on erroneous Transition games (Contrast-estimate:X0.112, F(1,53.275) = 7.191, P = .010). This suggests that making correct responses on Transition Revaluation games involved, to some degree, updating incorrect SR predictions, in the direction of correct MB predictions.

Although our primary interest was in investigating Transition Revaluation games, where SR and MB make divergent predictions, for purposes of comparison to prior literature, we also investigated Reward Revaluation games. On these games both MB and SR make the same predictions, and both predict correct revaluation. Prior studies examining similar manipulations have found greater predictive neural activity aligned with either MB or SR on correct responses, and thus contrast this to MF which predicts incorrect revaluations and no predictive neural activity (Doll et al., 2015; Wise et al., 2021). Although we observed directionally that SR alignment on these games was greater during correct compared to incorrect responses, this difference was not significant (Contrast-estimate: 0.015645, F(1, 98.514) = 0.12128, P = 0.7283). This null result may suggest that, on Reward Revaluation games, subjects were able to re-compute reward values prior to evaluation time (for example during Retraining, (Liu, Mattar, Behrens, Daw, & Dolan, 2021), see Discussion).

As with our behavioral results, all of the decoding results are robust to defining revaluation success using a continuous measure (Momennejad et al., 2017) as well as to using other strategies for controlling for Retraining acquisition failures, which remove fewer games (Supplementary Fig. 2 and 3).

Finally, because the foregoing analyses read out state predictions via category-specific activity in visual cortex, which may be elicited by other multi-modal association areas, they do not speak directly to which brain networks might encode the underlying predictive relationships. Previous theoretical studies have hypothesized hippocampus and medial prefrontal cortex (mPFC), to be candidate regions for predictive map utilized by the successor representation (Corneil & Gerstner, 2015; de Cothi & Barry, 2020; Geerts et al., 2020; Madarasz & Behrens, 2019; Russek et al., 2017; Stachenfeld et al., 2017; Yu, Park, Sweigart, Boorman, & Nassar, 2021). Although prior human neuroimaging studies have identified these areas as containing predictive representations (Barron et al., 2020; Boorman et al., 2016; Garvert et al., 2017; Howard, Gottfried, Tobler, & Kahnt, 2015; Wimmer & Büchel, 2019), because these studies only used a single step of state prediction, and did not include transition changes, these studies were not able to disambiguate predictions made by MB and SR. Investigating representational similarity in hippocampus and mPFC during the Observation Block, we identified predictive neural activity related to behavior, however, such predictive neural activity was consistent with either MB or SR state predictions, and did not adjudicate between them (Supplementary Fig. 4). Thus, a full determination of whether such activity reflects MB or SR predictions remains open for future work.

## Discussion

Although it is widely believed that the brain simulates the consequences of potential choices to evaluate them, we know very little about the mechanisms by which it manages to do so efficiently. Whereas traditional formulations of model-based choice evaluation postulate that sequences of action consequences be simulated one step at a time, this process can become computationally prohibitive (Daw & Dayan, 2014). In contrast, recent proposals have suggested that the brain may reduce the need for such time costly stimulation by relying on a temporal abstraction strategy, the successor representation (SR), whereby multiple steps of state predictions are learned from direct experience and used to replace the need for one-step simulation (Dayan, 1993; de Cothi et al., 2020; Geerts et al., 2020; Momennejad et al., 2017; Russek et al., 2017; Stachenfeld et al., 2017). Here, we provide novel neural evidence for this approach by demonstrating predictive neural representations that are both consistent with the state predictions used by the SR and predict corresponding evaluation behavior.

We accomplished this by using a task in which the SR produces behavioral errors following targeted changes to the task’s transition structure. Our key result is that these behavioral errors are accompanied by neural signatures of corresponding errors in multistep predictive representations that, according to the SR, give rise to the mistakes. This relationship was identified in prospective neural activity in sensory cortex during state evaluation. Notably, in one of our experimental conditions, Transition Revaluation games, the SR makes distinct predictions about behavioral errors from model-based evaluation. We observed that SR-predicted erroneous neural activity was accompanied by such SR-predicted, but not MB-predicted, behavioral errors. Altogether, the data are consistent with our hypothesis that underlying the behavioral errors during multi-step choice evaluation is a strategy of temporal abstraction: storing predictions of future states aggregated over multiple steps.

Our findings directly address two key gaps in the literature on predictive neural activity and choice evaluation. A previous study identified SR-consistent behavioral errors using a revaluation manipulation similar to the one we used here (Momennejad et al., 2017). However, the key inference about underlying neural representations made in that study, that these errors result from misprediction of future states, had not been verified. By identifying that such behavioral errors are accompanied by neural markers of SR-aligned incorrect state predictions, we provide positive evidence supporting that inference. Conversely, multiple previous studies had identified neural markers of predictive activity across numerous brain areas (Barron et al., 2020; Boorman et al., 2016; Deuker, Bellmund, Navarro Schröder, & Doeller, 2016; Doll et al., 2015; Howard et al., 2015; Stachenfeld et al., 2017; Wimmer & Büchel, 2019; Wise et al., 2021). However, such studies had not provided evidence that such predictive activity relate to behavioral choice evaluation in a manner consistent with SR evaluation. By demonstrating that predictive neural activity relates to variation in behavior across trials, and more specifically that this variation matches key predictions made by the SR, our findings thus provide key evidence that SR-aligned predictive neural activity reflects a mechanism for SR-aligned behavioral choice evaluation.

The key measure of predictive representations used here was prospective neural activity decoded from sensory cortex. Previous studies have identified that this measure reflects simulated outcomes of choices and relates to between subject variation in succeeding in reward revaluation (Doll et al., 2015; Wise et al., 2021). However, because these studies either required only a single of step state prediction or did not introduce any transition changes, they could not provide evidence distinguishing the key difference between SR and MB representations: multiple-step vs. one-step predictions, respectively. Additionally, unlike the current study, which utilized within-subject variation, these previous studies examined between-subject variation. Here, by utilizing a task requiring two steps of state prediction, and manipulating transitions in retraining, we were able, game by game, to dissociate temporally abstract multistep predictions (as in the SR) from the predictions that would be constructed by chaining together multiple one-step associations (as in MB).

Our ability to make this dissociation between MB-aligned versus SR-aligned state predictions relied on reasoning about which final state each algorithm would predict to follow either start state. In particular, on Transition Revaluation games, whereas an SR learned from the preceding trials would predict the outdated incorrect final state, MB would predict activation of the newly correct final state. Importantly, this reasoning is only valid under the assumption that MB made predictions with the correctly updated one-step model (because this is necessary for it to infer the correct two-step predictions). In particular, an MB process that failed to update its one-step transition model in the Retraining phase, would also produce incorrect final state predictions, along with incorrect revaluation behavior. Importantly, this alternative interpretation was refuted with data from behavioral valuations from each middle state (Supplementary Fig. 1) as well as final state category decoding from each middle state (Supplementary Fig. 3).

Related to this, a limitation of fMRI is that its slow temporal resolution does not allow examination of the temporal dynamics with which the divergent state predictions consistent with either SR or MB occur. However, future work, using faster timescale methods such as MEG (Liu et al., 2021; Wise et al., 2021), or possibly even novel approaches for fMRI (Schuck & Niv, 2019), may be able to further probe the dynamics of state representation, in order directly to test the prediction that any MB-consistent predictions are built up progressively by iterating steps one by one, while SR-consistent predictions are directly retrieved.

Another relevant design feature of this study is that subjects did not choose actions as they would in a typical sequential decision task, but rather evaluated states whose transition dynamics they had observed passively. In our previous work we measured human behavior in similar conditions both when the structures were passively learned and when they were learned following free choices (Momennejad et al., 2017). We observed that humans make analogous SR-predicted behavioral errors both in the passive framing, as in the present study, and in the more standard free choice framing. Although we would not expect key valuation processes to differ between these two framings, future work can examine neural markers of predictive representations in tasks where learning proceeds following free choices.

Whereas we were successful in identifying clear evidence for SR state prediction during evaluation from decoded activity in sensory cortex, we were unsuccessful at identifying the same signatures from the hippocampus and medial PFC following evaluation, using representational similarity analysis searchlights (Supplementary Fig. 4). In these areas, we identified that markers of predictive representations change coherently depending on revaluation behavior. However, these changes are consistent both with models that include SR as well as those that include only MB, and more detailed analyses to contrast the two did not produce conclusive results. We suggest that this might be due to lack of power – notably, unlike the decoding analysis, this analysis was not constrained by a separate localizer scan and thus required extensive correction for multiple comparisons. It is also possible, however, that changes in predictive representations (and various re-computation) may have occurred between the timepoint of evaluation (when the unambiguous SR-consistent results were measured using MVPA) and the later RSA measurement. We discuss one possibility for re-computation below.

It is important to stress that while we have reported affirmative evidence that SR is among the strategies employed, our results also show that the brain does not rely exclusively on the SR. Behaviorally, although participants made errors on Transition Revaluation games as predicted by the SR, they did not always do so: indeed, they were often able to succeed at correctly changing their preferences. Such successful revaluations were accompanied by neural markers of state predictions shifting away from incorrect SR predictions and toward correctly updated state predictions. This successful behavioral revaluation, along with the corresponding change in state prediction, implies that valuations and state predictions were, during those games, produced by MB simulation or replay: i.e., combining the one-step transition experiences from Training and Retraining phases, rather than relying on an SR learned directly from experienced multistep trajectories. Importantly, the MB process might be enacted by a variety of approaches that iterate one-step transition predictions to arrive at valuations. These approaches vary in whether the content of these one-step predictions reflect statistical averages or replayed memories from individual episodes, as well as what intermediate representations are computed en route to reward prediction. For example, whereas some approaches to valuation construct reward predictions directly from simulation or replay (e.g. “Dyna-Q” (Sutton, 1991) and basic MB value iteration), other recent approaches use either simulation or replay to update an SR and construct reward predictions from this (e.g. “SR-MB”, “SR-Dyna” (Momennejad et al., 2017; Russek et al., 2017)).

Despite the MB valuation implied by behavioral revaluation, and our observation of a change in neural state predictions between correct and incorrect revaluation games, we did not observe significant markers of MB-aligned state predictions on either correct Transition Revaluation games or on correct Reward Revaluation games. This may be because, for all the approaches to MB simulation or replay mentioned above, the timing with which such simulation or replay occurs is unconstrained. Although our study was designed to detect simulation occurring during ratings, it has been previously identified that MB simulation can occur prior to stimulus valuation, either during feedback or during rest periods between feedback and choice (Liu et al., 2021; Mattar & Daw, 2018; Momennejad et al., 2018). Thus, it is possible that (on correct trials) such simulation did occur, however it occurred prior to the rating phase during which we would have been able to measure it. This possibility suggests further research into understanding how the brain selects the time with which to re-compute reward predictions.

Despite our failure to observe direct neural markers of MB prediction, our key positive result is the observation of significant neural alignment with erroneous SR predictions during behavioral errors on Transition Revaluation games. These erroneous predictive representations were not observed when participants displayed accurate behavior. Both observations are consistent with our key hypothesis that out-of-date SR predictions were consulted during erroneous behavioral ratings, and that these predictions were either updated prior to, or not consulted during, correct ratings. Altogether this provides further evidence for the SR as a process for how the brain simplifies computations required for planning in multi-step tasks.

We conclude by pointing to two broader areas of research to which our result contributes. The first concerns the understanding of the computational and neural processes underlying behavioral inflexibility. In particular, the results provide further evidence that habit-like inflexible decisions can result from re-use of multi-step state predictions. Such state-prediction based cognitive habits may prove useful in understanding psychiatric conditions such as anxiety and depression, which are characterized by aberrant habit-like thought patterns as well as the development of idiosyncratic pessimistic predictions of future events (e.g. depressive schemas, (Beck, 1970; Huys, Daw, & Dayan, 2015)). Although it does not on its own explain why such idiosyncratic multistep state predictions develop in the first place, the SR suggests why they are difficult to change. In particular, updating the SR requires staging of choices followed by their multi-step consequences in time, and experience may take a long time to propagate over multiple steps in more complicated environments.

Finally, this work adds an additional thread to the multiplicity of process models for how the brain simplifies computations required for planning in multi-step tasks. Specifically, the SR joins other approximation strategies including pruning (Huys, Lally, et al., 2015), habitual selection of goals (Cushman & Morris, 2015), prioritized memory simulation (Mattar & Daw, 2018; Momennejad et al., 2018), action chunking and transfer of multi-step policies (Dezfouli & Balleine, 2013; Xia & Collins, 2021), and intermixing steps of model-based evaluation and model-free caching (Keramati, Smittenaar, Dolan, & Dayan, 2016). Understanding how these multiple processes interact, and how the brain might solve the meta-reasoning problem of deciding between multiple approximation strategies, presents a challenge and opportunity for future work.

## Methods

### Subjects

40 participants (23 female, mean age = 22.98, s.d. = 9.4) were recruited from the Princeton community. No statistical tests were used to plan this sample size, but this sample size was chosen, prior to data collection, to be above that typically used in the field. Subjects were paid $50 in addition to a performance-dependent bonus that ranged between $0 and $8. Princeton University’s ethics committee approved the study. All participants signed an informed consent form, reported no history of mental illness, and had normal or corrected-to-normal vision.

### Tasks

#### Functional localizer tasks

Subjects completed two functional localizer tasks, each consisting of a single scanner run lasting seven minutes. In the first task, the “block localizer”, subjects viewed 28-s image blocks followed by 12-s rest blocks. In each image block, 14 images of a single category (either faces, scenes or objects) were presented in randomized order for 1.4 seconds with a 600-ms inter-stimulus interval. Each category of image block recurred four times. Subjects were instructed to attend to the image on the screen and respond by button press as to whether the face, scene or object in the image was old or young. The images that made up the first localizer run were distinct from images used in the revaluation task.

In the second localizer task, the “event-based localizer”, in each of three blocks, subjects viewed 40 images, consisting of the first and last image of each sequence used in the task, presented in random order for 1-s with a 1-s to 3-s inter-trial interval, drawn randomly from a uniform distribution. Subjects were instructed to pay attention to each image. As a check for attention, fifteen times, randomly distributed through the task, subjects were presented with four images, and were asked to select the image they were most recently presented.

#### Reinforcement learning task

We used a reinforcement learning task, similar to experiment 1 in (Momennejad et al., 2017), with a few modifications to facilitate fMRI scanning (Fig. 1). Participants completed 10 games, each corresponding to one of three conditions: Reward Revaluation (four games), Transition Revaluation (four games) and Control (two games). Conditions were randomly interleaved for each participant.

In a Training phase, subjects passively experienced two sequences, each presented six times in random, interleaved order (Fig. 1a). During training, each image was presented for 1.5-s with an inter-image interval of 250-ms and an inter-sequence interval of 750ms. The final image of either sequence was paired with either a high or low reward, signified by an image of $100 or $1 respectively, and the first two images were paired with a blurred dollar image to indicate that they were not rewarding. Additionally, each image had a background color that prescribed its position in the sequence (Fig. 1b). The sequences were arranged so that the first two images of either sequence were from an object category and the final image of either sequence was a face or scene respectively. Whether a face or scene was paired with the high or low reward was randomly counterbalanced. Each sequence was shown 6 times, randomly interleaved. While an image was on the screen, subjects were required to press a button to indicate whether the person, object or scene in the image was young or old. After every third sequence, subjects were randomly shown either the last images or the middle images of either sequence and asked to select the image they thought would more likely lead them to the high reward.

Following Training, in the Rating 1 phase, subjects were presented with the first two images of either sequence, one at a time and in random order, constrained so that the first images of either sequence were presented before the middle images of either sequence. Subjects used button presses to position a slider bar so as to rate each image in terms of how much they would like to start a new sequence from that image. Each image presentation was TR-locked. Images were on the screen for 4 seconds, during which time subjects provided a rating by moving a slider bar. Following each rating was a randomly selected 8-,10-, or 12-second interval.

Subjects then completed a Retraining phase similar to the Training phase, except that each sequence started from its middle state instead of from the start state (so that only the middle image and final image were experienced) (Fig. 1a). Additionally, depending on the condition, subjects were exposed to a change in each sequence. In Reward Revaluation games, the rewards paired with either sequence’s final image were switched. In Transition Revaluation games, the final image that followed the middle image for either sequence were switched. In Control games, no change occurred. Each partial sequence was presented three times, in randomly interleaved order. After every second sequence position, subjects were randomly shown either the last images or the middle images of either sequence and asked to select which image they thought would more likely lead the high reward.

Following retraining, in a second rating phase (Rating 2), subjects rated the first two images of either sequence, i.e., the start states and middle states, in the same order as the first rating (Fig 1a). Again, they were instructed to rate their preference for starting a new sequence from the presented state. Each Rating was followed by a randomly selected 8-, 10-, or 12-second inter-rating interval.

Finally, following the Rate 2 phase, in an Observation Block phase, subjects viewed a sequence of 30 images, consisting of the first and third image of either sequence. Images were presented in a random order, subject to the constraint that the four transitions defined by the first image of either sequence being immediately followed by the third image of either sequence each occurred at least once. Each image was presented for 1 second, with a 1-3 second inter-trial interval. As an attention check, randomly interspaced in the sequence were four probes, in which the four images in the Observation Block appeared on the screen and subjects had to select the image they most recently observed.

The task was explained to participants using a cover story in which each game corresponded to visiting a new city and that in each city, each day they would take photographs of each location that they visited along a travel path. Participants were instructed that for each path, the path’s final location depended on the middle location and that the middle location depended on the start location. Participants were instructed that at each location they would take a photograph (which they saw on the screen) and that some of the photographs also earned them money. While a photo was on the screen, subjects were required to categorize whether the subject in the photo was old or young. Participants were told that while they were in a city, both the value of certain photographs as well as parts of paths themselves could change. For the rating phase, participants were told to rate how much they would like to visit a location (cued by that location’s photograph) and then complete a path starting from that location.

### Removal of games

For all analysis presented in the main text, in order to limit our analysis to games where MB, SR and MF make clear predictions, we restricted our analysis to games in which (a) subjects’ ratings for start and middle states (during Rating 1) reflected a correct preference for the higher valued state presented in Training, (b) subjects’ ratings for middle states (during Rating 2) reflected the newly correct preference for these states, and (c) subjects did not provide the same rating in Rating 2 to each start image. In short, these criteria ensured that we analyzed games in which participants had both accurately learned the value of different trajectories to begin with, and that they had accurately learned the change in either rewards or transitions in every game we analyzed. Across all subjects, this resulted in removing 39 games due to (a), an additional 77 games due to (b) and an additional 35 games due to (c). Removing these games caused one subject to be removed from the dataset entirely, leaving 39 subjects for analysis.

Removing games based on (a) was necessary because correct versus incorrect revaluation only has meaning when the baseline prior to the change occurring was successfully learned. For example, if an MF agent somehow acquired incorrect preferences prior to the change in rewards or transitions, even it could then produce correct ratings in Rating 2 (since it would not need to change its preferences at all). Removing games based on (b) was necessary because, on Transition Revaluation games, MB and SR only make distinct predictions about Rating start states when both MB and SR have successfully updated their values of middle states. For example, if an MB agent failed to update its one-step transition model (as would be reflected by failing to update preferences for Rating 2 middle states), then it would also fail to update preferences for Rating 2 start states and predict identical behavior to SR and MF. Lastly, removing games based on (c) is necessary in order to define Rating 2 in each game as being discretely correct or incorrect.

In the supplementary materials, we explore an alternative way of controlling for retraining acquisition failures. Additionally, we use a continuous-valued measure of revaluation success. Thus, for this analysis we did not exclude games based on (b) and (c). As a result, for this analysis only 35 games were removed and no subjects were removed entirely.

### Behavioral analysis

For the analysis presented in Fig. 2, for each game, we categorized each Rating 2 phase as being correct or incorrect depending on whether the subject rated the newly higher-valued start state greater than the newly lower valued start state. The successor representation predicts that subjects will make more errors on Transition Revaluation games compared to Reward Revaluation games. We tested this prediction by fitting a generalized linear mixed effects model (with a logistic response distribution) to predict a game’s correctness (dummy coded as 1 for correct and 0 for incorrect), where the explanatory variables contained one dummy-coded variable for each condition (Reward Revaluation, Transition Revaluation and Control). Here and elsewhere we use mixed effects models with full random effects for subjects on all parameters, so as to correctly capture the repeated measure structure of the experiment and data, which are sometimes unbalanced across cells. In order to compare the proportion of correct responses between conditions, we compute a contrast between fixed effects for each condition using MATLAB’s “coef-test” which computes the significance test on the F-statistic of the contrast (here, with infinite degrees of freedom). This function returns fixed and random effect estimates, *β*, which reflect logit-transformed log-odds correct for each condition. In order to covert these into measurements of group and participant-specific proportion correct for each condition, which is plotted in Fig. 2c, it is necessary to pass each estimate, *β*, through an inverse logit function, 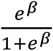.

### fMRI procedures

Gradient echo T2*-weighted echo-planar images (EPI) with blood oxygen level dependent (BOLD) contrast were collected on a 3-T Siemens Skyra MRI scanner. Slices were collected using a multi-band sequence with interleaved slice acquisition with an acceleration factor of 2. Seventy axial slices (2×2×2 mm) were acquired in oblique orientation of 30 degrees to the anterior-commissure posterior-commissure line with repetition time (TR) of 2s and TE of 30 ms, 90 degree flip angle, 192 mm field of view. Following the main task, a high-resolution T1-weighted anatomical image (magnetization-prepared rapid-acquisition gradient echo sequence 1×1×1 mm voxels) was also collected. In order to construct a field map, we also collected two EPIs with opposite phase-encoding polarity, a TR of 9.240s, TE of 79ms and otherwise identical parameters to the task images.

FSL topup (Smith et al., 2004) was used to construct a fieldmap from the phase-encode-reversed pairs. Preprocessing was then conducted using SPM12 (Wellcome Trust Centre for Neuroimaging, http://www.fil.ion.ucl.ac.uk/spm/). SPM’s Field MAP toolbox was used to warp functional images and realign for head motion. Functional images were then co-registered across runs and to the structural image and resampled to 2×2×2 mm voxels. Functional images smoothed with a 5-mm FWHM Gaussian kernel for the decoding analysis and 2-mm FWHM Gaussian kernel for representational similarity analysis. Decoding analysis was performed in native space. For representational similarity analysis, images were normalized to Montreal Neurological Institute (MNI) space.

### Decoding Prospective Neural Activity During Evaluation

#### Regions of Interest

To assist our classifier in defining good features, we restricted training data to BOLD signal from anatomical areas known to be selective for faces and scenes. We created a single mask, by combining anatomical masks, defined in (Julian, Fedorenko, Webster, & Kanwisher, 2012), corresponding to face selective regions (Fusiform Face Area, Occipital Face Area and Superior Temporal Sulcus) and scene selective regions (Parahippocampal Place Area, Retrosplinial Cortex, and Transverse Occipital Sulcus). For each subject, this mask was warped to native space using SPM 12.

#### Decoding

We used data from the two localizer tasks to train a three-way L2 normalized, logistic regression classifier (penalty = 1100) to predict which image category the subject was viewing. The classifier was implemented using the Princeton Multi-Voxel Pattern Analysis Toolbox (https://github.com/PrincetonUniversity/princeton-mvpa-toolbox). We defined features for each image presentation by extracting the BOLD signal from the anatomical masks, temporally filtering (using spm_filter, RT = 2, H = 128) and z-scoring for each run. Data from the TR closest to 2 TRs following image onset was then used as features for that image presentation, whose label corresponded to either face, scene or object. The penalty parameter as well as how many TRs to offset features from image presentation were selected as those that maximized cross validation performance at classifying a single held out image presentation from the event-related task.

To examine predictive neural activity during Rating 2, during each rating (when an image of an object was on the screen) the trained classifier was applied to the TR corresponding to 2 TRs after the (TR locked) onset of the rated image. The classifier provided a readout of the (inverselogit) probability that the BOLD signal (temporally filtered and z-scored for each run) during this TR reflected activity scene when viewing a face, scene or object category. For each game, we computed a score measuring the correctness of the category predictions implied by the decoded neural activity. Specifically, if, following the Retraining phase image A would be expected to be followed by a face and image B would be expected to be followed by a scene, Category Prediction Correctness was computed as

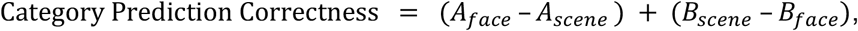

where *A_face_* is the classifier readout of face probability during onset of image A. Note that, for the Transition Revaluation condition, this measure was taken with respect to correct integration of change implied by the Retraining phase.

For the analysis presented in Figure 3c and d, we fit a linear mixed effects model to estimate this score across all games (and across all subjects). The model contained one dummy coded explanatory variable for each Revaluation condition (Reward Revaluation, Transition Revaluation, Control) which indicated whether the game to which the classifier measure corresponded was from that Revaluation condition. Additionally, for each Revaluation condition, it also included an explanatory variable for that condition, interacted with whether behavior for that game was correct or incorrect. Specifically, for each Revaluation condition, this variable was 0 if the game was not from that condition. If the game was from that condition, this variable was 1 if behavior on that game was correct and −1 if behavior was incorrect. MATLAB’s “coef-test”, which computes the significance test on the F-statistic of the contrast (with degrees of freedom estimated using Satterthwaite approximation), was used to compute significance of contrasts between fixed effects of this model.

## Acknowledgements

We wish to thank Oliver Vikbladh and Aaron Bornstein for helpful conversations, and Nicholas DePinto and Leigh Nystrom for assistance with scanning protocols. This project was supported by a NIH grant 1R01MH109177, part of the CRCNS program, as well as a NIH NRSA grant 5F31MH110111.

## Supplementary Materials

### Robustness of behavioral and decoding results

Here, in order to further explore the robustness of our results, we present an additional analysis of the behavioral as well as the decoding results using a continuous measure of the degree to which subjects updated valuations following each game’s Retraining phase. Specifically, rather than compute a discrete measure of correct or incorrect revaluation, for each game, we compute a revaluation score, which reflects preference for the newly correct image in Rating 2 (Image 2 – Image 1), subtracting relative preference for that image in Rating 1 to control for any differences in the magnitude of initial acquisition (Gershman et al., 2014; Momennejad et al., 2018, 2017). Formally, revaluation score is computed as follows:

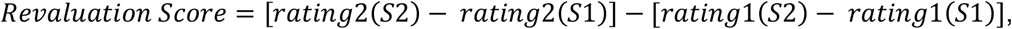

where *rating 2*(*S1*), refers to the second rating of the start state from the initially non-rewarding sequence (Fig. 1a).

Because, compared to the discrete measure of correct versus incorrect, the revaluation score measure provides a finer measure of degree of learning, this measure affords other means of controlling for reward change and transition change acquisition failures. Thus, for this analysis, we do not remove games where the subject failed to demonstrate correct preference for re-valued middle images in Rating 2. We also do not remove games where subjects provided an equal rating for both images. We still remove trials, however, where subject’s valuations during Rating 1 did not reflect the correct ordering of which image was more valuable. Altogether, this results in removal of 35 games with no subjects removed entirely.

Supplementary Figure 1 shows that subjects had lower revaluation scores on Transition Revaluation games compared to Reward Revaluation games (in agreement with Fig 2c; Contrast estimate = −0.253, F(1,37.570): 15.813, P < .001). Subjects also had higher revaluation scores on Reward Revaluation games compared to Control games (Contrast estimate = −0.576, F(1, 41.912) = 84.889, P < .001) as well as on Transition Revaluation games compared to Control games (Contrast estimate = −0.323, F(1, 34.428) = 34.428, P < .001).

**Supplementary Figure 1:**
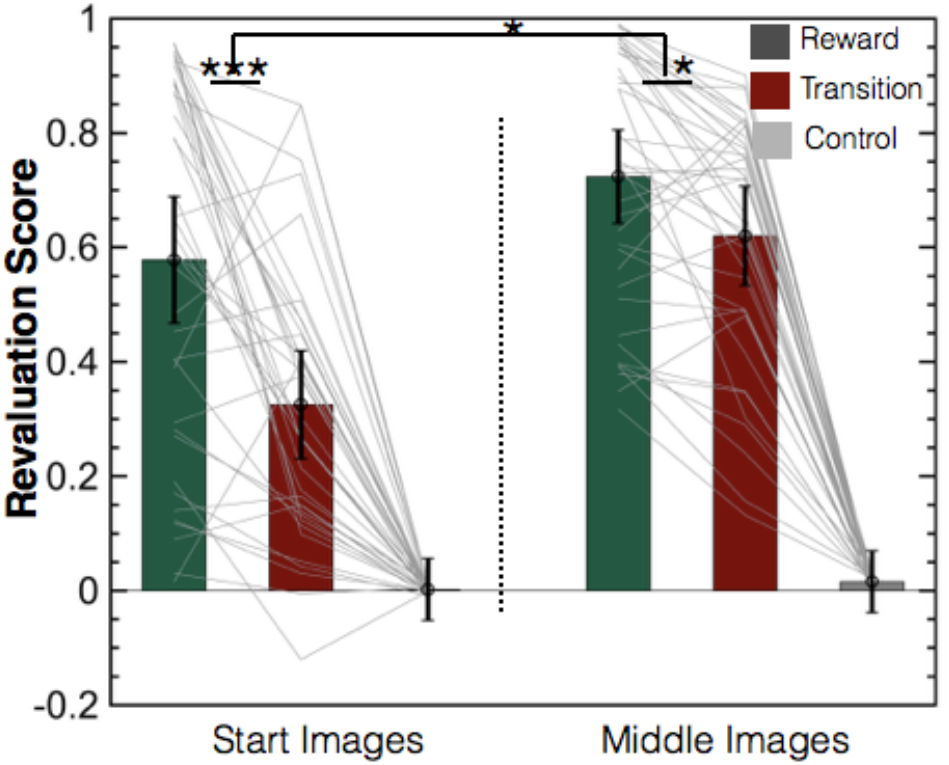
Revaluation score measurements for each condition for both start and middle images (N = 40 subjects). Bar heights reflect measured revaluation scores based on group-level, fixed effect estimates from a linear mixed effects model relating revaluation score to condition and sequence position (start or middle). Error bars show 95% confidence intervals. Grey lines show fitted subject-level random effects estimates. ***P < .001, *P< .05, linear mixed effects model.

Although a difference in revaluation errors between Reward Revaluation and Transition Revaluation games is the signature prediction of the SR, such a difference could also arise even using only MB, if the new associations were imperfectly learned during Retraining, and in particular if transition changes were less successfully acquired than reward changes (e.g. due to a lower learning rate for updating transition vs reward estimates). Thus, it is important to directly assess and control for the success of the underlying re-learning. For this analysis, we did this by repeating the same analysis, but this time measuring change in relative preference for the middle image of either sequence (Image 4 – Image 3, Fig. 1a), which are the images whose associations were directly re-experienced during the Retraining phases. Any differential success at acquiring reward vs transition changes should be visible as a difference in these scores between conditions; but, importantly, any such difference could only give rise to a difference in scores of equal size at the start images.

Instead, although there was a difference in revaluation scores at the middle images between Reward Revaluation and Transition Revaluation games (Contrast estimate = 0.104, F(1, 40.671) = 5.801, P = .021), this difference was smaller than the difference in revaluation scores for starting images (Contrast estimate = 0.149, F(1,6.341) = 6.342, P = .015). This indicates that poorer acquisition of transition changes relative to reward changes could not account for participants’ selective impairment in revaluation for transition games.

For these analysis related to Supplementary Figure 1, in order to compute differences in revaluation score between conditions, we fit a linear mixed effects model predicting revaluation score with a dummy-coded explanatory for each combination of condition (reward revaluation, transition revaluation or control) and sequence position (start or middle), again with per-subject random effects for each predictor. In order to take contrasts between conditions, here and for each following analysis using linear mixed effects models, we used MATLAB’s “coef-test” function, estimating degrees of freedom using the Satterthwaite method.

Supplementary Figure 2 shows that fMRI decoding results also remain the same using the revaluation score measure. When revaluation scores on Transition Revaluation games were 0, suggestive of SR behavior, Category Prediction Correctness was less then 0 (Estimate = −1.514, F(1,45.33) = 12.216, P = .001), consistent with SR neural predictions. We additionally observed a positive relationship between Category Prediction Correctness and revaluation score on Transition Revaluation games (Contrast estimate = −0.215, F(1,39.646) =5.726, P = .022), suggesting that as revaluation scores become more correct, suggestive of MB, decoded category predictions move in the direction of MB predicted activity. As with the discrete measure, we did not observe a relationship between Category Prediction Correctness and revaluation score on Reward Revaluation games (Contrast estimate = 0.001, F(1, 69.912) = 0.000, P = .987).

**Supplementary Figure 2:**
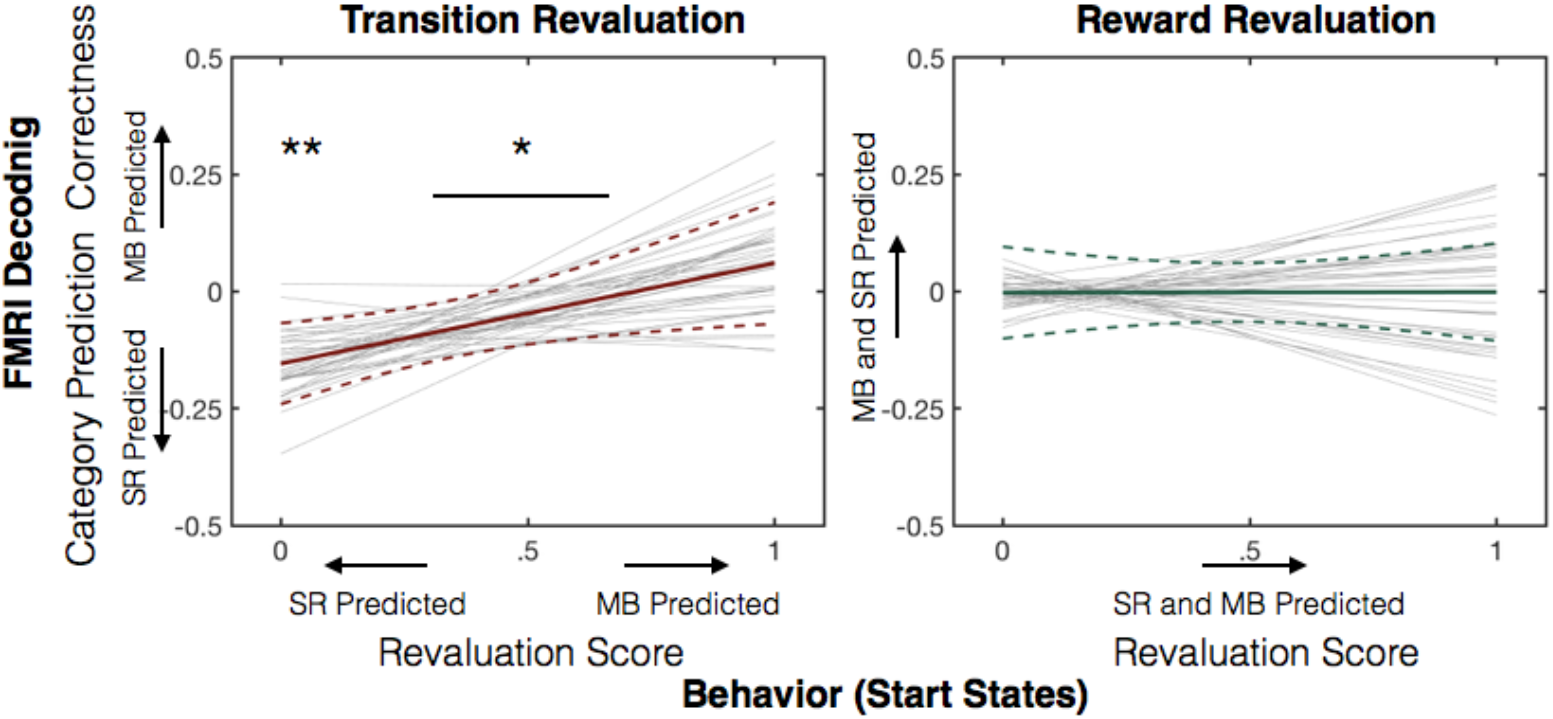
Relationship between Category Prediction Correctness and revaluation score on Transition Revaluation and Reward Revaluation Games (N = 40 Subjects). Solid colored lines show fixed effects group prediction from linear mixed effects model predicting Category Prediction Correctness. Dashed lines show 95% confidence intervals. Grey lines show subject-level random effects predictions. ** P < .005, *P < .05, linear mixed effects model.

Using revaluation score measure behavioral correctness and not removing incorrect revaluations of middle states also enabled us to conduct an additional analysis that further controlled for the degree of correct acquisition of the underlying contingencies. In particular we were able to measure predictive neural activity during ratings for the middle states (the ones directly updated during Retraining). If the relationship between incorrect predictive neural activity elicited by start states and errors were due to an MB system that failed to acquire the new contingencies, and not the SR, we would expect that revaluation errors for start states would be explained by greater erroneous final state predictions from the middle states.

To examine this possibility, we also measured predictive neural activity from the middle states on the Rating 2 of Transition Revaluation games. From this, we measured Category Prediction Correctness from each middle state, in addition to each start state. For each middle state, this measure was computed equivalently to how it was computed for start states. Specifically, it was the relative activity for the correct minus incorrect (post Retraining phase) final image category. We treated this measure of Category Prediction Correctness, from each middle state, as a measure of transition-change acquisition. We then examined whether a relationship between failures of transition-change acquisition (measured as more negative Category Prediction Correctness from each middle image) and low start state revaluation scores could account for the relationship between more negative Category Prediction Correctness from the start state and low start state revaluation scores. Supplementary Figure 3 shows that the relationship between this measure and start state revaluation scores was smaller than the relationship between Category Prediction Correctness from the start states and revaluation scores, thus ruling out MB with acquisition failures as an explanation for our findings (Contrast estimate: 0.272, F(1, 42.496) = 5.348, P = .026).

**Supplementary Figure 3:**
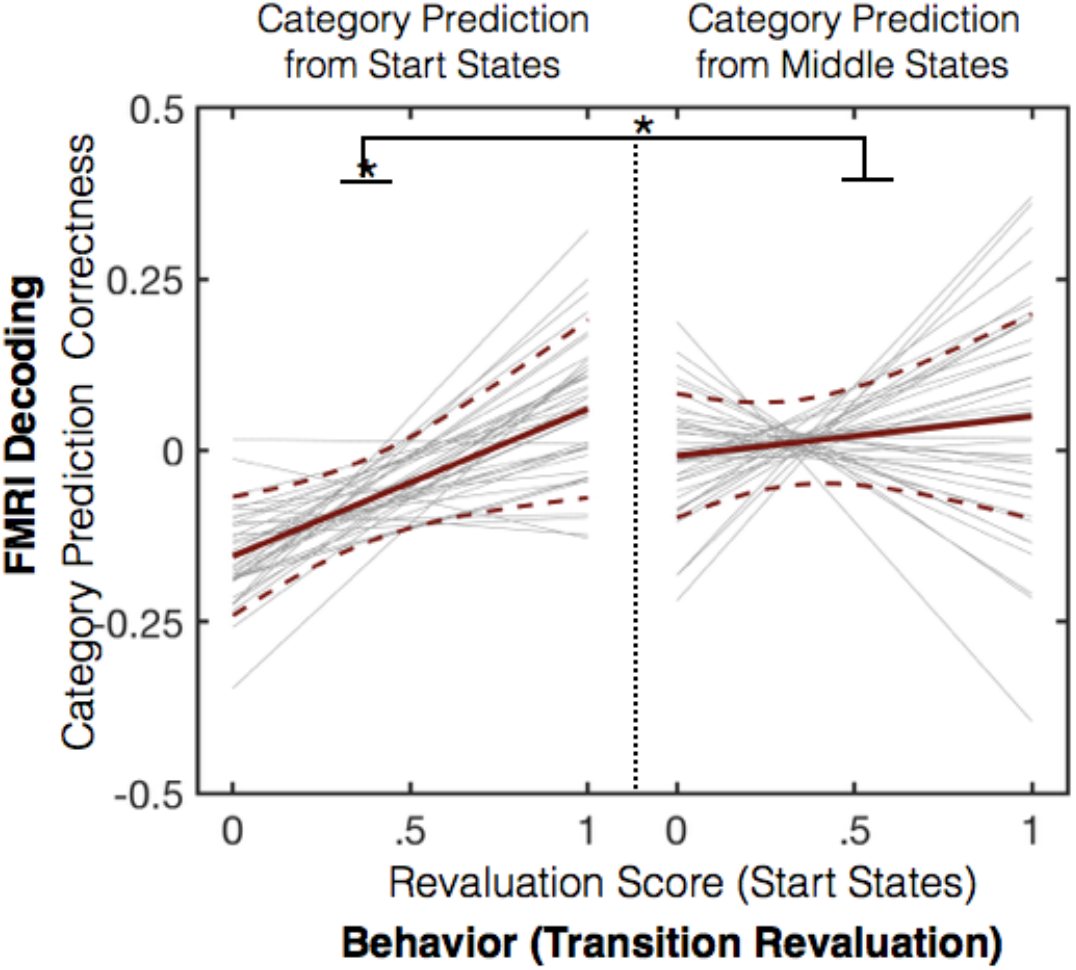
Relationship between relative Category Prediction Correctness and start state revaluation score for measured category predictions during ratings of start states versus middle states. (Transition Revaluation games; N = 40 Subjects). Solid colored lines show fixed effects group prediction from linear mixed effects model predicting relative erroneous final image category evidence. Dashed lines show +/− 95% confidence intervals. Grey lines show subject-level random effects predictions. *P < .05, linear mixed effects model.

For the analysis in Supplementary Figures 2 and 3, we fit a linear mixed effects model to predict Category Prediction Correctness for both Start and Middle state ratings across all games, across all subjects. The explanatory variables consisted of a dummy-coded explanatory variable for each combination of condition (Reward Revaluation, Transition Revaluation or Control) and sequence-position (start or middle). For example, an explanatory variable corresponding to start state ratings from the Reward Revaluation condition contained a 1 if the rating was of a start state and was done on a Reward Revaluation game. It contained a 0 if the rating either was of a middle state, or corresponded to a Transition Revaluation or Control game. For each of these explanatory variables, an additional variable was also included which interacted these condition variables with the revaluation score for the start state for that game.

### Representational Similarity Analysis Results

Previous theoretical studies have hypothesized that a predictive map underlying SR multi-step state predictions exists in hippocampus and medial prefrontal cortex (mPFC) (Corneil & Gerstner, 2015; de Cothi & Barry, 2020; Geerts et al., 2020; Madarasz & Behrens, 2019; Russek et al., 2017; Stachenfeld et al., 2017; Yu et al., 2021). Some previous studies have identified representations in these areas that contain predictive representations (Brunec & Momennejad, 2020; Garvert et al., 2017; Howard et al., 2015; Schapiro, Turk-Browne, Norman, & Botvinick, 2016; Wimmer & Büchel, 2019) with some relations to successful reward revaluation (in the form of sensory preconditioning, (Barron et al., 2020)). However, because these studies only used a single step of state prediction, and did not introduce transition changes, they were unable to examine whether those predictive representations reflect predictions made by SR or MB. In order to investigate whether these representations encode SR predictions, we applied representational similarity analysis (RSA; (Kriegeskorte, Mur, & Bandettini, 2008)) to FMRI data collected during the Observation Block phase of each game (Supplmentary Fig. 4). Because RSA is not tied to image categories, (which are associated with specific areas of higher visual cortex), it could be used to index state representations in other parts of the brain, and thus allowed investigation of representations in hippocampus and mPFC.

**Supplementary Figure 4:**
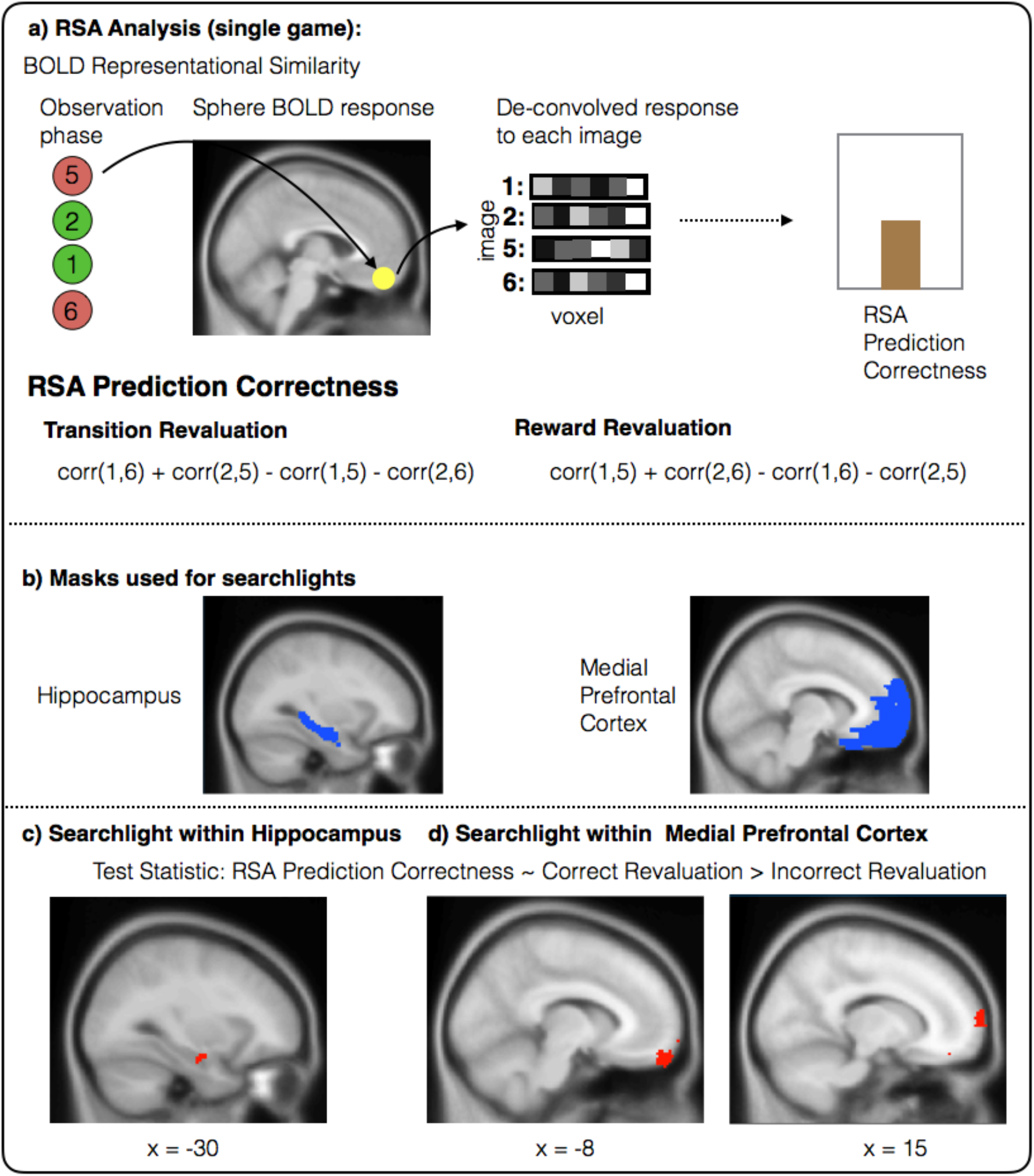
Representational Similarity Analysis. **a) Structure of RSA analysis.** Within each 3-voxel-radius sphere, we modeled the response to each of the four images (corresponding to the first and last state of either sequence) presented during the Observation Block for each voxel within a sphere. We then computed correlations between the responses to the start image of either sequence and the last image of either sequence, and treat the correlation as a measure of the prediction that the given final image would follow the given start image. RSA Prediction Correctness measure the extent to which these predictions are correct following the Retraining phase. Specifically, on Reward revaluation games, they measure the extent to which state 5 is predicted to follow state 1 (corr(1,5); subtracting the opposite) and the extent to which state 6 is predicted to follow state 2. On Transition revaluation games, following the change in one-step transition predictions, they measure the extent to which state 6 is predicted to follow state 1 and state 5 is expected to follow state 2. **b) Masks used for searchlight analysis. c,d) Results from each searchlight measuring how RSA Prediction Correctness changes depending on whether revaluation behavior is correct or incorrect.** Note that, as explained in the text, this analysis can provide evidence for relationship between predictive representations in these areas and behavior however, does not disambiguate whether or not those predictive representations reflect SR. **c) Results from bilateral hippocampus searchlight**. One cluster in left anterior hippocampus displayed a significant interaction of condition and Rating-2-Correctness (Peak MNI = [−32 −8 −16], P_SVC_ < .025). **d) Results from medial PFC searchlight.** One cluster in left orbitofrontal cortex (left; Peak MNI: [2, 62, −18], P_SVC_ < 0.025) and one in right medial prefrontal cortex (right; Peak MNI [14, 72, 10], P_SVC_ < 0.025) showed displayed a significant interaction of condition and Rating-2-Correctness. c,d) Image threshold P_uncorrected_ < .005), Images reflect RSA results applied to 39 subjects.

We ran separate searchlight analyses within a mask for each of these regions (Schapiro et al., 2016) (Fig 4b). For every 3-mm radius sphere, centered on each voxel within the mask, we computed the correlation in the BOLD response to each start state’s image and each final state’s image. After correcting this measure for the same measure of similarity from data during an initial pre-task baseline exposure (to eliminate any baseline image similarity effects not due to task learning), we treated this difference as a readout of the predictive representation encoding the multi-step prediction that a particular final state would follow a particular start state (Supplementary Fig 4a). From these correlations, we defined a score, RSA Prediction Correctness, analogous to the score used in the decoding analysis. In particular, on Reward Revaluation games, RSA Prediction Correctness reflected the degree to which image 1 was expected to be followed by image 5 (as indexed by higher correlation) and the extent to which image 2 was expected to be followed by image 6. In contrast, on Transition Revaluation games, following the one-step transition change, correctness reflected the degree to which image 1 would be expected to be followed by image 6 and 2 would be expected to be followed by image 5.

As with the decoding analysis, the key difference between MB and SR predictions occurs on Transition revaluation games, where, following the Retraining phase, SR makes incorrect multistep predictions (as learned during Training), but MB makes correct multi-step predictions which have been re-computed using an updated one-step transition model. Our key prediction was that Transition Revaluation errors, predicted by SR to occur behaviorally, would co-occur with SR predicted representations, reflecting incorrect multi-step predictions, and as observed in the decoding analysis. However, examining in hippocampus and mPFC, isolated to erroneous Transition revaluation games, did not reveal RSA Prediction Correctness to be significantly different from 0 in either direction, suggesting that Transition errors could not be attributed to either incorrect, SR predicted neural similarity, or correct, MB predicted similarity. Additionally, no effect was observed in either direction, in either mask, on correct Transition revaluation games, suggesting that, on their own, these games also could not be attributed to either MB or SR predictive activity.

We next examined whether there was a difference in RSA Prediction Correctness between correct and incorrect Transition Revaluation games, as was also observed in the decoding analysis. Importantly, this difference, in absence of the negative main effect on erroneous Transition Revaluation games that was observed in decoding analysis, would not, on its own, disambiguate whether incorrect responses are due to SR or MF. Rather, this variation could be explained by MB responsibility for correct responses and correct state predictions trading off with either MF responsibility for incorrect responses and incorrect predictions (zero predictive activity), or SR responsibility for incorrect responses and incorrect predictions (incorrect predictive activity).

However, we did not observe a difference in Correct RSA Alignment between correct and incorrect Transition revaluation games in either mask. Altogether this suggests that representational similarity in hippocampus and mPFC could not be attributed to predictions of either SR or MB, and that no change in balance between these occurred dependent on behavior.

We next considered a weaker measure of whether predictive representations in these areas relate to behavior, in ways that do not disambiguate between MB and SR. Previous studies, using manipulations akin to Reward Revaluation, with only a single step of state prediction, have demonstrated that neural markers of state prediction reflect predictions that are more correct when revaluation behavior is more correct. This has been demonstrated in both visual cortex (Doll et al., 2015) as well as in hippocampus and medial PFC (Barron et al., 2020). We investigated whether this effect exists in representational similarity in Hippocampus or medial PFC by aggregating across Reward Revaluation games and Transition Revaluation games and examining whether neural similarity reflected more correct multi-step state predictions on games where revaluation was successful compared to when it was not successful. Across all revaluation games, we identified clusters in both masks where RSA Prediction Correctness was greater on correct compared to incorrect revaluation games (Supplementary Fig. 4c; **Anterior Hippocampus**: 9 voxels, P_SVC_ < .025; Peak MNI = [−32 −8 −16], Contrast-estimate = −0.873, F(1,117.56) = 15.865, P_uncorrected_ < 1e-4; Supplementary Fig. 4d; **left orbitofrontal cortex**: 65 voxels; P_SVC_ < 0.025; Peak MNI: [2, 62, −18], Contrast-Estimate = −1.1668, F(1,87.39) = 16.073, P_uncorrected_ < 1e-4; **right mPFC**: 53 voxels, P_SVC_ < .025; Peak MNI: [14, 72, 10], Estimate: −1.3972 (SE = 0.312), F(1, 158.43) = 20.024, P_uncorrected_ < 1e-4) .

Importantly, this effect is predicted both by models in which MB trades off with MF and SR as well as those where MB trades off with MF alone. In addition to variation performance on Transition Revaluation games, which described above, in absence of a main negative effect on incorrect Transition Revaluation games, is not discriminative of SR influence, variation in Reward Revaluation games could be explained by MF responsibility for errors and reduced RSA Prediction Correctness and either MB or SR responsibility for correct responses and increased RSA Prediction Correctness. Thus, although this effect demonstrates that measures of predictive representations change depending on correctness, it is consistent with predictive representations reflecting either MB alone, or a combination of MB and SR.

We also examined this effect isolated to Reward Revaluation games. Here we found significant effects in both masks (**left Orbitofrontal Cortex**: 1: 209 voxels, P_SVC_ < 0.025, Peak MNI: [−10, 56, −16], P_uncorrected_ < 1e-5; 2: 69 voxels, P_SVC_ < 0.025, Peak MNI: [4, 44, −20], P_uncorrected_ < 1e-4; **left Hippocampus**: 7 voxels, P_SVC_ < 0.025, Peak MNI: [−30, −22, −10], P_uncorrected_ < 1e-3). This effect is surprising, given the null effect of behavior on Category Prediction Correctness in the decoding analysis.

Altogether, these results suggest discrepancies in predictive representations as measured from representational similarity analysis and those elicited during evaluation, as measured by decoding from visual cortex. We briefly discuss two potential reasons for this discrepancy. One reason for the failure to observe effects in the RSA, which were observed during sensory cortex during evaluation, may simply be a lack of power. Whereas for the decoding measurement we had a mechanism of identifying, a priori, the measurement corresponding to predictive neural activity, for representational similarity, we were forced to use a searchlight approach, and thus correct for hypothesis over many individual tests. Alternatively, it is possible that predictive representations observed during choice in sensory cortex, and those observed after choice in medial PFC and hippocampus, do not reflect identical predictions. As discussed in the Discussion section of the paper, this could occur if multi-step state predictions had been re-computed sometime following the evaluation yet prior to the RSA measurement. Finally, it is possible that the brain contains multiple predictive maps, in different locations, and that that these can become out of sync from one another, or alternatively can be differentially updated at different times.

### Representational Similarity Analysis Methods

Based on a priori hypotheses, we defined two volumes: one covering medial prefrontal cortex (Wake Forest Pickatlas, (Maldjian, Laurienti, Kraft, & Burdette, 2003), Brodmann areas 9 and 10, dilated 5 voxels and limited to x in [−25, 25], MNI coordinates), and one covering bilateral hippocampus (conjunction of left and right Automatic Anatomical Labeling defined hippocampus masks). For each volume, using the Princeton MVPA Toolbox, we ran a “searchlight” (Kriegeskorte, Goebel, & Bandettini, 2006). For each voxel in a given volume, we defined a spherical ROI (searchlight) of 6-mm radius centered on that voxel, excluding voxels outside the mask. For each spherical ROI, we computed a statistic measuring the correctness of the predictions implied by the correlations between the first and third image in either sequence. For Reward Revaluation games, this score was computed as

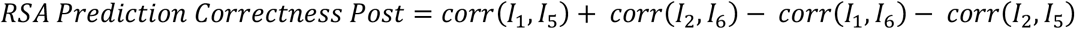

where *I*_1_ and *I_2_* are the vectors of responses of each voxel to the first image (state) of either sequence respectively (as presented in training; Fig. 1), *I*_5_ and *I*_6_ are the vectors of responses of each voxel to the last image (state) of either sequence respectively, and corr is the Fisher z-transformed Pearson correlation.

For Transition Revaluation games, following the change in one-step Transitions in the Retraining phase, this score was computed as:

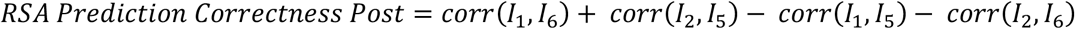

The input to these correlations was a map over voxels within each sphere of responses to each image, with hemodynamic effects deconvolved. We estimated this deconvolution for each image by constructing a general linear model in SPM, with a single condition for each image with box car regressors for when the image was on the screen. Events from other phases (not the Observation Block phase) of the trial including the category of the image on the screen, whether the subject was making a rating, whether the subject was making a choice during the training or retraining phase, and the value of any money image on the screen were included as nuisance regressors. Each condition was convolved with a standard hemodynamic response function. We also modeled each equivalent image as it was presented in the event-based localizer task; and computed a score for similarity in between those images as well, *RSA Prediction Correctness Pre*. This score was identical to *RSA Prediction Correctness Post*, except that it was computed using image presentations from the event-based localizer task. We computed our test statistic of interest by subtracting the score from the event based localizer from the score from the Observation Block, *RSA Prediction Correctness = RSA Prediction Correctness Post — RSA Prediction Correctness Pre*. The purpose of this subtraction was to correct for any similarity between task images caused by visual features.

For each searchlight, we computed *RSA Prediction Correctnesss* for each game included in our decoding analysis, across all subjects. We then fit identical linear mixed effects models to this score as we did for the decoding analysis. For each analysis, uncorrected p-values were computed for every voxel evaluating the significance of fixed (group-level average) effects.

Within each mask, P-values were then corrected for multiple comparisons using small volume correction for the volume over which the search took place. We estimated smoothness parameters by providing the map of the square-root of model deviance for the model fit in each voxel. Because all contrasts were taken with respect to a single model, this map was the same for all presented contrasts. This map was passed to AFNI’s 3dFWHMx, which was used to compute parameters to estimate smoothness (using the newer ACF method) (Cox, 1996). These parameters were used by AFNI’s 3DClustSim to estimate the null distribution over cluster sizes given the cluster-defining threshold of p < .001 uncorrected. For each mask, we computed the number of voxels needed for a significance level of p < .025 (so as to Bonferroni correct for additional family-wise error due to the test of two ROIs). This revealed a threshold of 6.7 voxels for the hippocampus mask and 44.5 voxels for the medial PFC mask.

